# Bidirectional crosstalk between epithelial-mesenchymal plasticity and IFNγ-induced PD-L1 expression promotes tumor progression

**DOI:** 10.1101/2022.02.03.478950

**Authors:** Gerhard A. Burger, Daphne N. Nesenberend, Carlijn M. Lems, Sander C. Hille, Joost B. Beltman

## Abstract

Epithelial-Mesenchymal Transition (EMT) and immunoevasion through upregulation of Programmed Death-Ligand 1 (PD-L1) are important drivers of cancer progression. While EMT has been proposed to facilitate PD-L1-mediated immunosuppression, the molecular mechanisms of their interaction remain obscure. Here we provide insight into these mechanisms by proposing a mathematical model that describes the crosstalk between EMT and Interferon gamma (IFNγ)-induced PD-L1 expression. Our model shows that via interaction with microRNA-200 (miR-200), the multistability of the EMT regulatory circuit is mirrored in the PD-L1 levels, which are further amplified by IFNγ stimulation. This IFNγ-mediated effect is most prominent for cells in a fully mesenchymal state, and less strong for those in an epithelial or partially mesenchymal state. Additionally, bi-directional crosstalk between miR-200 and PD-L1 implies that IFNγ stimulation allows cells to undergo EMT for lower amounts of inducing signal, and that IFNγ presence accelerates EMT and decelerates Mesenchymal-Epithelial Transition (MET). Overall, our model agrees with published findings and provides insight into possible mechanisms behind EMT-mediated immune-evasion; and primary, adaptive, or acquired resistance to immunotherapy. Our model can be used as a starting point to explore additional crosstalk mechanisms, as an improved understanding of these mechanisms is indispensable for developing better diagnostic and therapeutic options for cancer patients.

**Graphical Abstract:** 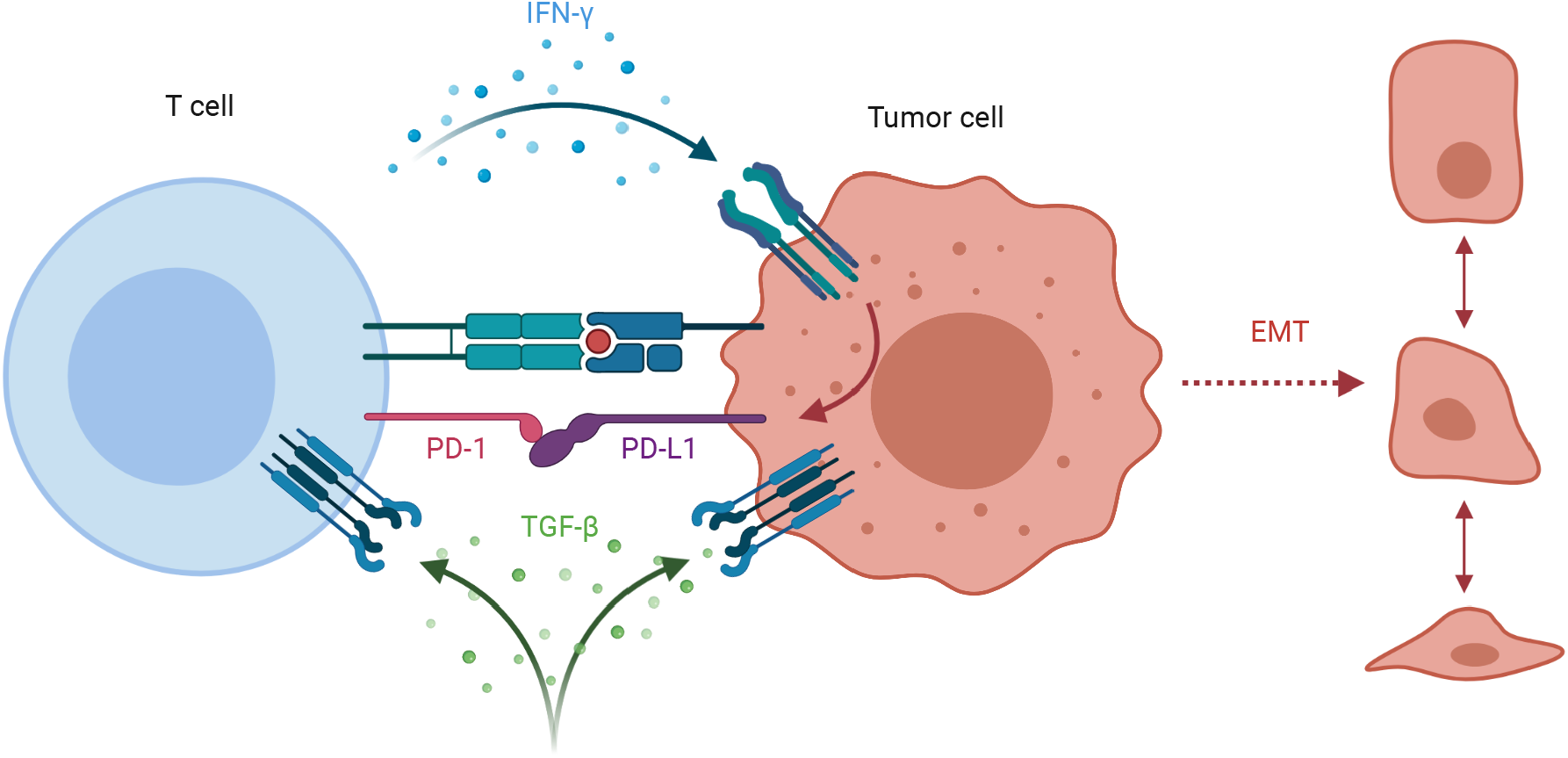

Schematic overview of the crosstalk between Epithelial-Mesenchymal Transition (EMT) and Interferon gamma (IFNγ)-induced Programmed Death-Ligand 1 (PD-L1) expression. IFNγ-induced PD-L1 expression promotes the occurrence of EMT. EMT, also induced by, for example, Transforming Growth Factor Beta (TGFβ), increases PD-L1 expression levels, facilitating tumor immunoevasion.

## Introduction

Activating invasion and metastasis and evading immune destruction are well-established hallmarks of cancer, complementary capabilities that enable tumor growth and metastatic dissemination (Hanahan and Weinberg, 2011). Because metastasis is the main cause of cancer mortality, a thorough understanding of the interaction between these hallmarks is essential for developing therapeutic approaches, yet they are often studied separately (Meirson, Gil-Henn, and Samson, 2020). Consequently, the interplay between metastatic dissemination and immune evasion remains poorly understood.

In recent years, Epithelial-Mesenchymal Plasticity (EMP), the ability of cells to interconvert between intermediate E/M phenotypes along the Epithelial-Mesenchymal Transition (EMT) spectrum (Jing Yang et al., 2020), has been extensively studied because of its crucial role in invasion and metastasis (reviewed in Derynck and Weinberg, 2019; Williams et al., 2019; W. Lu and Kang, 2019). Moreover, EMP is associated with therapeutic resistance (reviewed in Staalduinen et al., 2018; Williams et al., 2019), including resistance to immunotherapy (reviewed in Terry et al., 2017). One of the mechanisms through which cancer cells acquire immune resistance is by activation of immune checkpoint pathways, which under physiological conditions are indispensable for self-tolerance and modulation of the immune response (Pardoll, 2012).

A frequently upregulated checkpoint protein in tumors is Programmed Death-Ligand 1 (PD-L1) (Okazaki and Honjo, 2006). Its corresponding receptor Programmed Death-1 (PD-1) is expressed on the cell membrane of T cells, and PD-1–PD-L1 interaction induces a variety of immunosuppressive effects, such as inhibition of T cell proliferation, survival, and effector functions (Zitvogel and Kroemer, 2012). Tumor cells can express PD-L1 by two general mechanisms: adaptive immune resistance, where PD-L1 expression is upregulated in response to inflammatory factors (such as Interferon gamma (IFNγ)) produced by an anti-tumor immune response, and innate immune resistance, where PD-L1 expression is upregulated in response to constitutive oncogenic signaling (Pardoll, 2012). Such constitutive signaling could, for example, be caused by EMT, and indeed several links between EMT and PD-L1-mediated immunoevasion have been reported in the literature (reviewed by Jiang and Zhan (2020), who conclude that additional mechanistic studies are urgently needed).

Here we study the post-transcriptional regulation of PD-L1 by the microRNA-200 (miR-200)–Zinc Finger E-Box Binding Homeobox 1 (ZEB1) axis, which is one of the proposed mechanisms underlying the interplay between EMT and immune resistance (Limo Chen et al., 2014; Noman et al., 2017; Martinez-Ciarpaglini et al., 2019). The binding of miR-200 to the mRNA of PD-L1 inhibits mRNA translation and stimulates miRNA decay (Mitarai et al., 2009; Baccarini et al., 2011). The miR-200-ZEB1 axis is “a motor of cellular plasticity” (S. Brabletz and T. Brabletz, 2010) and is considered to be part of the EMT core regulatory network (Nieto et al., 2016). Various mathematical models of EMT that include the miR-200-ZEB1 axis have been developed, which have contributed to a better mechanistic understanding of EMT (Jolly, Tripathi, Somarelli, et al., 2017; Burger, Danen, and Beltman, 2017; Jing Yang et al., 2020). Recently, Sahoo et al. (2021) presented a mathematical model interconnecting a minimal EMT network and PD-L1 using shifted Hill functions to investigate immune-evasive strategies of hybrid E/M states, assuming an indirect effect of PD-L1 on EMT. Model analysis showed that both the stable hybrid and full EMT phenotypes resulting from the model were associated with high PD-L1 levels. However, although EMT scores determined from gene expression levels across a large panel of cell lines indeed exhibited a pattern of gradually increasing PD-L1 expression with increasing EMT score, there was no clear dichotomy as expected from the mathematical model. This difference may be explained by the indirect nature of the feedback of PD-L1 to EMT as implemented in the model of Sahoo et al. (2021). Moreover, the impact of adaptive, IFNγ-driven PD-L1 expression, i.e., the primary reason for PD-L1 upregulation (Mühlbauer et al., 2006; S. Chen et al., 2019), was not taken into account in their model.

To describe the full interactions expected between adaptive and innate immune resistance and to study the impact of direct, mutual feedback between EMT and PD-L1, we connected a “core” EMT model (the simplified Ternary Chimera Switch (TCS) model (Jolly, Tripathi, D. Jia, et al., 2016)) to a model for IFNγ-induced PD-L1 expression, which we developed based on an extension of a published JAK–STAT model (Quaiser, Dittrich, et al., 2011). Combining these two models was achieved by adding a mutually inhibitory feedback loop between miR-200 and PD-L1, which we described using appropriate miRNA-mRNA dynamics (M. Lu, Jolly, Gomoto, et al., 2013). Analysis of our model shows that IFNγ-induced PD-L1 expression is expected to greatly accelerate EMT and decelerate the reverse Mesenchymal-Epithelial Transition (MET) process. Moreover, IFNγ-induced PD-L1 lowers the required level of EMT-inducing signal, leading to an overall larger probability of EMT in tumors with a high IFNγ expression compared to tumors with a low expression. Vice versa, and consistent with Sahoo et al. (2021), a full EMT induced via other signals greatly upregulates PD-L1 expression, which IFNγ further amplifies. However, in our model, the hybrid EMT phenotype only moderately affects PD-L1 expression, which is due to the mutual feedback between EMT and PD-L1. Finally, we show that our model findings are broadly consistent with published experimental results in an extensive comparison to experimental data. Overall, our analysis illustrates how crosstalk between EMP and IFNγ-induced PD-L1 production can result in immune evasion and contribute to invasion and metastasis.

## Results

### Modeling IFNγ-induced PD-L1 expression and EMT

To study the interplay between IFNγ-induced PD-L1 expression and EMT, we first need a quantitative description of these major processes. Although IFNγ-induced PD-L1 expression via the JAK–STAT pathway (Fig. 1A) has been extensively studied (Garcia-Diaz et al., 2017; Ivashkiv, 2018), to the best of our knowledge, no mathematical model of this entire process exists. Existing models focus on IFNγ-induced JAK–STAT signaling (reviewed in Gambin et al., 2013) and usually do not take into account direct targets of Signal Transducer and Activator of Transcription (STAT) such as Interferon Regulatory Factors (IRFs) and Interferon-Stimulated Genes (ISGs). To model IFNγ-induced PD-L1 expression, we used the simplified JAK–STAT model by Quaiser and Mönnigmann (2009) (Fig. 1B, left box), which is a truncated version of a more detailed model by Yamada et al. (2003). In this model, STAT1p_2 reaches its maximum at about 30 minutes after IFNγ exposure (Fig. 1C, left), which represents a much shorter time scale than the time scales at which PD-L1 expression or EMT is expected to occur. Therefore, we simplified this model further by describing the steady state of its output, i.e., phosphorylated Signal Transducer and Activator of Transcription 1 (STAT1) homodimer (STAT1p_2), with a Gompertz function (Supplementary Methods). We then extended this model with the production of IRF1 mediated by STAT1, and production of PD-L1 mediated by IRF1 (Fig. 1B, middle box, and Fig. S1A). We modeled both of these processes with shifted Hill functions, which led to realistic dynamics (Rateitschak et al., 2010) in which PD-L1 becomes expressed on a time scale of hours (Ghosh et al., 2021) (Fig. 1C, middle, and Fig. S1B).

**Figure 1:**
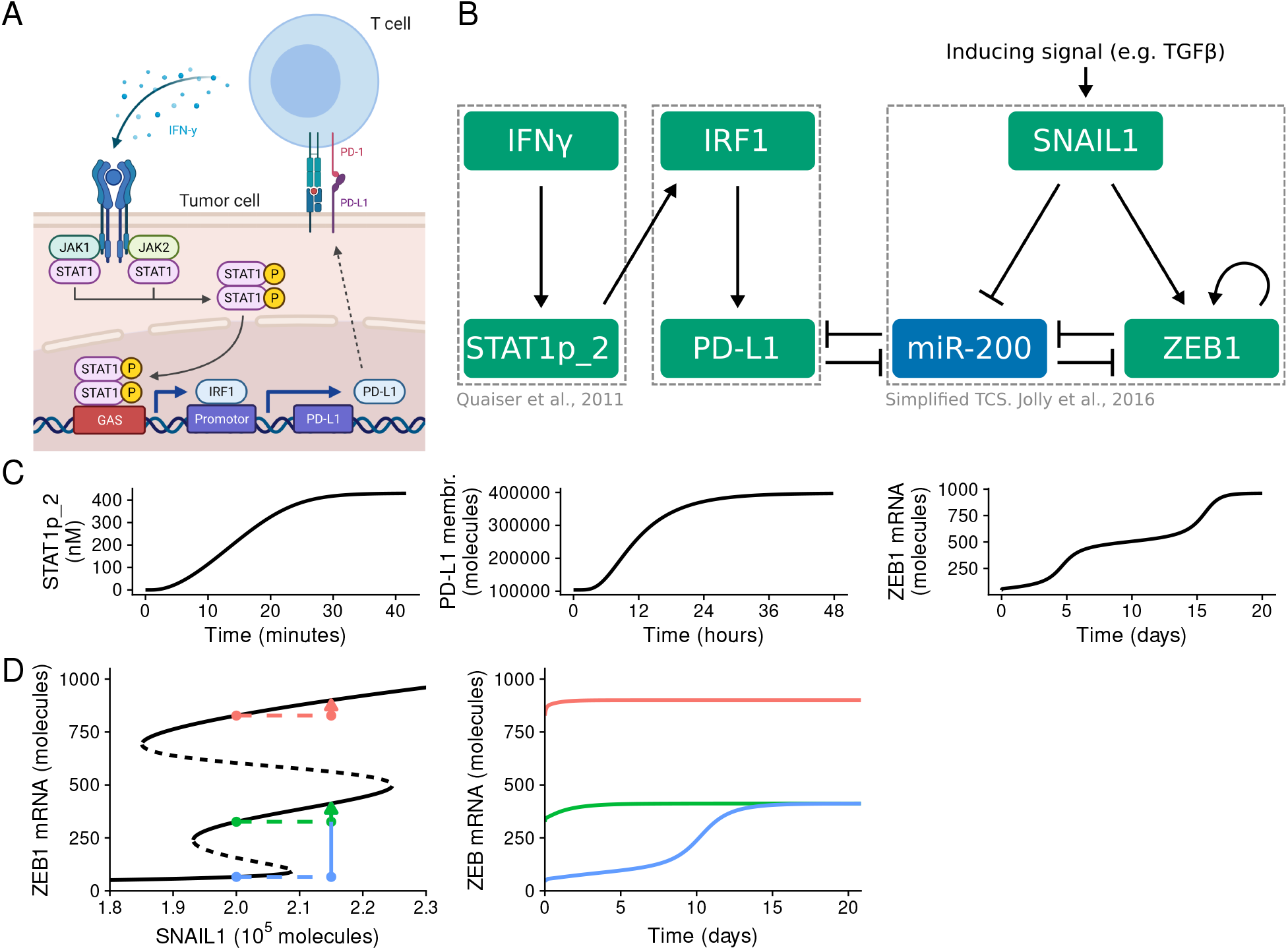
PD-L1 and EMT dynamics occur on distinct time scales. (A) Schematic diagram of the signaling pathway regulating IFNγ-induced PD-L1 expression. (B) Schematic diagram of the full model combining IFNγ-induced STAT signaling (left box), PD-L1 expression (middle box), and the simplified TCS model for EMT dynamics (right box). (C) Temporal dynamics of separate models (boxes in B) without connection to the other parts. This includes IFNγ-induced STAT1 signaling (left) with IFNγ = 0.1 nm, STAT1-induced PD-L1 expression (middle) with STAT1p_2 = 430 nm, and SNAIL1-induced EMT (right) with SNAIL1 = 2.3 × 10^5^ molecules. (D) Bifurcation diagram illustrating how EMT phenotype depends on the inducing signal in the TCS model (left, black lines), and how the eventual phenotype also depends on the initial state of the system, illustrated by temporal dynamics after a sudden increase of SNAIL1 from 2.0 × 10^5^ molecules to 2.15 × 10^5^ molecules given three different initial states (colored lines).

For EMT, various mathematical models have already been developed (Jolly, Tripathi, Somarelli, et al., 2017; Burger, Danen, and Beltman, 2017). We elected to use the TCS model developed by Jolly, Tripathi, D. Jia, et al. (2016) (Fig. 1B, right box), which is a simplified model compared to prior work (M. Lu, Jolly, Levine, et al., 2013) and readily extensible. Notably, the TCS model exhibits three stable states, representing an epithelial phenotype (E), an intermediate phenotype (E/M), and a mesenchymal phenotype (M) (Fig. 1D, left, black lines). Consistent with the results for the full TCS model (cf. Fig. 6 in M. Lu, Jolly, Levine, et al., 2013), a full EMT in the simplified TCS model takes approximately twenty days, with a slowing down of the transitioning to the mesenchymal phenotype around ZEB1 expression levels for the hybrid phenotype (Fig. 1C, right). Both the initial state of the system and the EMT-inducing signal determine the phenotype that the system attains in the long run (Fig. 1D, colored lines). Moreover, it is important to note the difference in time scales for the dynamics of the different model components: whereas STAT signaling and PD-L1 expression occur on a time scale of minutes and hours, a full EMT transition from E to M occurs on a time scale of days (Fig. 1C).

### IFNγ amplifies the increase in PD-L1 caused by EMT primarily for mesenchymal cells

Because we aimed to study the impact of IFNγ-mediated PD-L1 on EMT and vice versa, we connected the separate model parts (Fig. 1B, boxes) by adding mutual inhibition between PD-L1 and miR-200 (Limo Chen et al., 2014; Alsuliman et al., 2015; Noman et al., 2017) using the theoretical framework for miRNA-TF dynamics by M. Lu, Jolly, Gomoto, et al. (2013). For this combined model (Fig. 1B), we studied how the system responds to different levels of SNAIL1 (activated via, for example, Transforming Growth Factor Beta (TGFβ)) and IFNγ (Fig. 2). In the absence of IFNγ, the previously reported tristable ZEB1 levels (M. Lu, Jolly, Levine, et al., 2013) translate to similar tristability in PD-L1 expression at the cell membrane (blue line in Fig. 2A). Here, the highest PD-L1 expression occurs for mesenchymal cells and the lowest for epithelial cells. Note that the PD-L1 expression of hybrid cells is only slightly increased compared to epithelial cells, a prediction that is different from the recent study by Sahoo et al. (2021). Notably, the model predicts that exposure of cells to IFNγ greatly amplifies PD-L1 expression for all phenotypes, thereby also amplifying the differences in PD-L1 expression between those EMT phenotypes (Fig. 2A, orange line; Fig. 2B). Nevertheless, in the presence of IFNγ the hybrid phenotype has a PD-L1 expression level similar to that of epithelial cells, and only mesenchymal cells are expected to have substantially higher PD-L1 levels.

**Figure 2:**
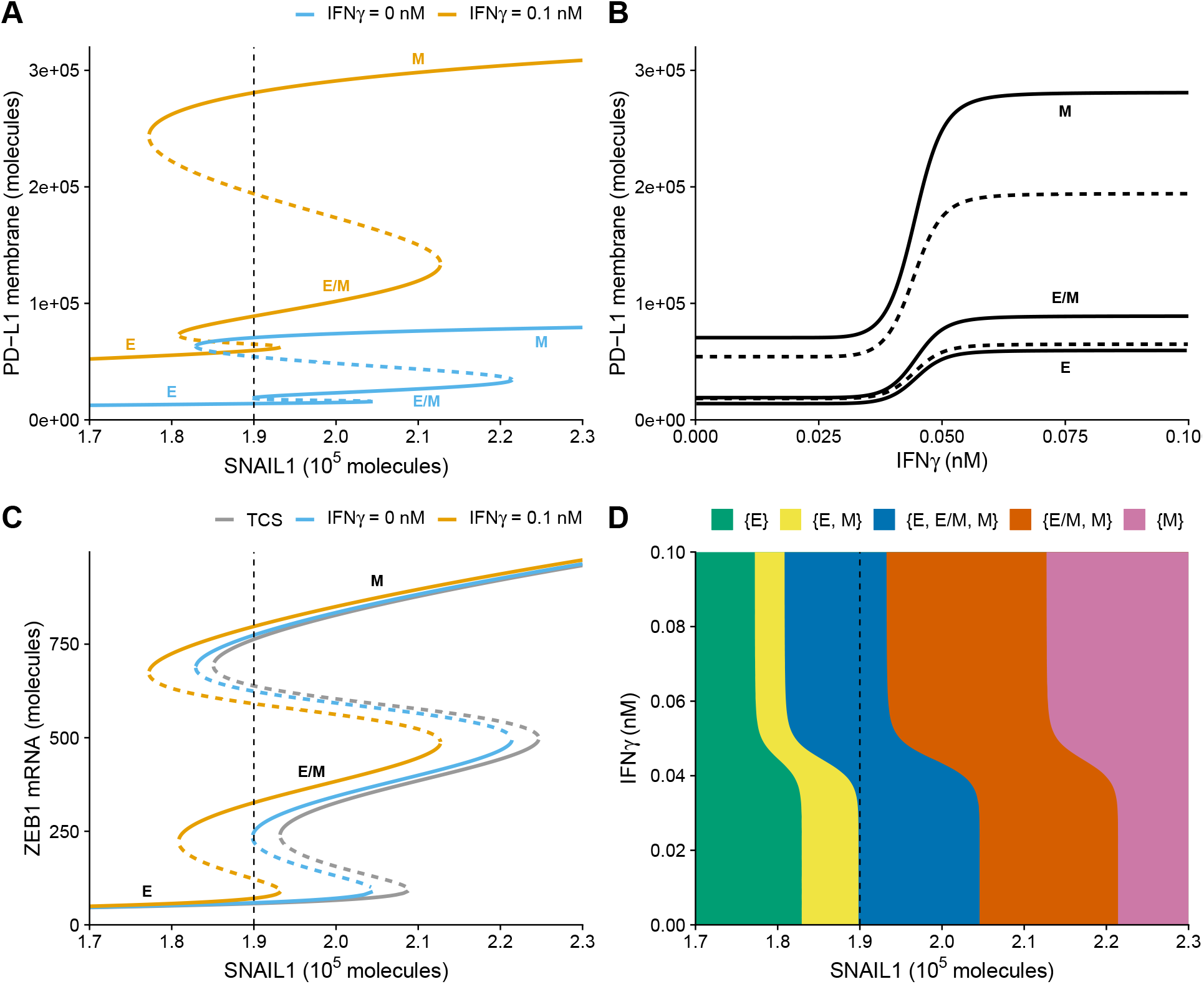
Model-predicted mutual influence of IFNγ-driven PD-L1 expression and EMT. (A)-(C) Bifurcation diagrams illustrating how the steady-state expression of PD-L1 on the membrane depends on SNAIL1 in the absence (blue) and presence (orange) of IFNγ (A), how the steady-state expression of PD-L1 on the membrane depends on IFNγ, considering a fixed SNAIL1 level of 1.9 × 10^5^ molecules (B), and how the steady state of ZEB11 mRNA depends on SNAIL1 in the TCS model (gray), and in our extended model in the absence (blue) or presence (orange) of IFNγ (C). Stable equilibria (representing E, E/M, and M phenotypes) are indicated by solid lines and unstable equilibria by dashed lines. (D) Phase diagram showing how the presence of stable equilibria (colored regions, indicated in legend) depends on IFNγ and SNAIL1. Vertical dashed lines in (A),(C), and (D) show the SNAIL1 concentration used in (B).

### IFNγ promotes the occurrence of EMT

We next asked what the impact of IFNγ is on ZEB1, i.e., a central EMT-regulating Transcription Factor (TF). Even in the absence of IFNγ, cells in our model can undergo EMT for lower levels of SNAIL1 compared to the original TCS model (leftward shift of the blue curve in Fig. 2C compared to the gray curve). This is because PD-L1 and miR-200 mutually influence each other, and the low basal expression of PD-L1 thus leads to a reduced amount of miR-200, in turn affecting EMT. In the presence of IFNγ, the bifurcation diagram shifts even further to the left (Fig. 2C, orange curve), which implies that according to our model, IFNγ induces EMT through PD-L1 upregulation. Note that in the presence of IFNγ, both partial and full EMT more readily occur than in its absence.

We created a phase diagram to provide an overview of how the stability of EMT phenotypes depend on IFNγ and SNAIL1 levels (Fig. 2D). In this phase diagram, the IFNγ-induced leftward shift is clearly visible. Additionally, it shows that the total range of SNAIL1 for which the hybrid E/M phenotype, a particularly aggressive EMT phenotype (Lüönd et al., 2021, and reviewed in Jolly, Somarelli, et al., 2018), can (co-)exist is unaffected.

To investigate the role of the included JAK–STAT regulation, we created a simplified crosstalk model where we replaced JAK–STAT signaling with a generic inducing signal *I* (Fig. S2A). The influence of this inducing signal is more gradual than the influence of IFNγ in the full model, yet the same leftward shift is present (Fig. S2B, cf. Fig. 2B), suggesting that IFNγ-induced JAK–STAT signaling makes the response highly “switch-like”. Additionally, we used a sensitivity analysis on parameters used in the negative-feedback loop between miR-200 and PD-L1, which showed that both the predicted leftward shift of SNAIL1 levels and the amplification of PD-L1 by IFNγ are robust to parameter change (Fig. S3). In conclusion, our model predicts that local presence of IFNγ promotes partial and full EMT by lowering the threshold of additional EMT-inducing signals.

### IFNγ-driven PD-L1 expression accelerates EMT

Besides the steady states for EMT status and PD-L1 expression that the combined regulatory network can reach in the long run, in practice, it will also matter how long it takes to reach these states. Therefore, we also investigated our combined regulatory network’s temporal dynamics and compared them to the temporal dynamics of the separate models.

To study the impact of PD-L1 expression on the temporal dynamics of an EMT transition and vice versa, we started with a stable epithelial phenotype (SNAIL1 = 1.7 × 10^5^ molecules) and simulated transition to a fully mesenchymal phenotype by a sudden increase of SNAIL1 (to 2.3 × 10^5^ molecules). In our combined model of PD-L1 expression and EMT with the double-negative feedback loop between miR-200 and PD-L1, several changes in the temporal dynamics of EMT can be observed relative to the simplified TCS model (blue lines in Fig. 3, left column): First, the decrease in miR-200 during EMT affects the PD-L1 mRNA and membrane PD-L1 (which are not present in the TCS model) at a similar time scale as miR-200. Thus, the effect of EMT on PD-L1 expression takes place on a much slower timescale than for PD-L1 expression driven by IFNγ (for comparison, see Fig. 1C, middle panel). Second, because of the double-negative feedback loop, the increase in PD-L1 over time speeds up the miR-200 decrease (compared to the TCS model without the feedback loop). Because of miR-200’s role as a critical suppressor of several EMT-TFs (Chung et al., 2016; Bracken, Scott, and Goodall, 2016), this accelerates the EMT process by about five days, mainly by reducing the time cells remain in a state close to the hybrid (E/M) phenotype before converting to a fully mesenchymal phenotype. Induction of PD-L1 by IFNγ enhances this accelerating effect by PD-L1, decreasing the time required for a full EMT by another five days, thus cutting the twenty days needed for a full EMT in the simplified TCS model in half (orange line in Fig. 3, left). Again, the PD-L1 expression level evolves slowly compared to the rapid increase observed on a time scale of hours for the model describing just IFNγ signaling (Fig. 1C). This difference in time scales of initial IFNγ-driven PD-L1 increase and of further PD-L1 increase due to EMT is especially apparent when considering a scenario where cells are simultaneously exposed to IFNγ and a TGFβ (Fig. S4, left column). In summary, the double-negative feedback loop between miR-200 and PD-L1 accelerates EMT, and IFNγ amplifies this effect.

**Figure 3:**
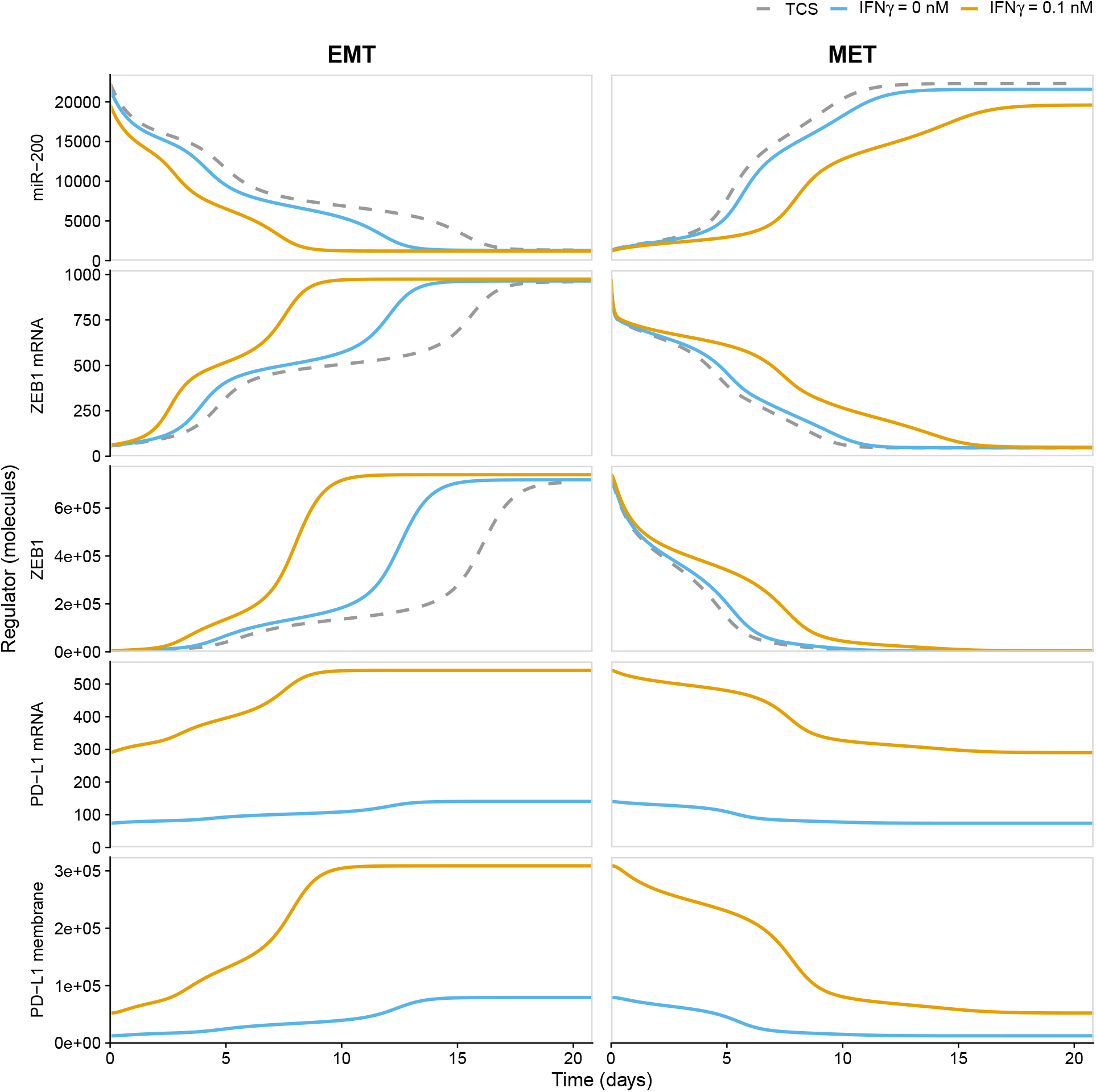
Temporal dynamics of EMP. EMT (left) and MET (right) for the simplified TCS model (dashed gray), and our combined model (Fig. 1B) with IFNγ = 0 nm (blue) and IFNγ = 0.1 nm (orange). For EMT, cells in an epithelial state with SNAIL1 = 1.7 × 10^5^ molecules undergo a full EMT by increasing SNAIL1 to 2.3 × 10^5^ molecules; for MET, SNAIL1 is decreased to the original value of 1.7 × 10^5^ molecules again.

### IFNγ-driven PD-L1 expression decelerates MET

Finally, we investigated whether similar changes occur in the temporal dynamics of the reverse process in which mesenchymal cells transition to an epithelial phenotype, i.e., MET. To this purpose, we started with a stable mesenchymal phenotype where SNAIL1 = 2.3 × 10^5^ molecules, which we instantaneously decreased to 1.7 × 10^5^ molecules. In the simplified TCS model, this SNAIL1 decrease led to a direct transition from the mesenchymal into the epithelial phenotype (gray dashed line in Fig. 3, right), i.e., without a substantial slowing down of the dynamics around a hybrid phenotype.

As anticipated from the observed EMT dynamics showing that the double-negative feedback loop between PD-L1 and miR-200 favors the mesenchymal phenotype, this loop also affects MET, albeit in the opposite direction, i.e., it decelerates MET. However, this effect is considerably smaller than the effect in the forward transition (blue lines in Fig. 3, right), which is likely the result of MET not transitioning through the hybrid state at a slow pace since the miR-200/PD-L1 interaction primarily accelerated EMT by shortening the time spent there (Fig. 3 left). The addition of IFNγ further decelerates MET, primarily due to the long-lasting suppression of miR-200 (orange lines in Fig. 3, right). Note that simultaneous lowering of SNAIL1 and removal of IFNγ do lead to an MET occurring at a similar speed as in the original TCS model (Fig. S4, right column), demonstrating the important role of IFNγ in the EMT dynamics. In conclusion, the PD-L1-miR-200 double-negative feedback loop is expected to slow down MET considerably, and high IFNγ concentrations strengthen this effect.

## Discussion & Conclusion

Here we have used a mathematical model to describe the crosstalk between IFNγ-induced PD-L1 expression and EMT. We showed that merely adding the reported interaction between PD-L1 and miR-200 gives rise to tristability in the PD-L1 levels, where a mesenchymal state corresponds with high PD-L1 expression and an epithelial state with low PD-L1 expression. The difference in PD-L1 levels between the stable EMT states is amplified by adding IFNγ stimulation. Additionally, we showed that adding this crosstalk reduces the amount of SNAIL1 required to undergo EMT. This reduction is dependent on the IFNγ concentration but is present even in the absence of IFNγ (i.e., compared to a model without the PD-L1-miR-200 crosstalk). Finally, we showed that this crosstalk accelerates the forward EMT process and decelerates the reverse MET process, and that IFNγ amplifies these effects.

To assess the extent to which the model results agree with recent studies, we have compiled a summary of published experimental reports on the link between EMT and PD-L1 and indicate which findings are qualitatively consistent with our results (Table 1). Note that we omitted several papers that propose different mechanisms underlying EMT/PD-L1 crosstalk and lacked experimental findings that could be compared to our simulations (Kumar et al., 2017; Bouillez et al., 2017; Suda et al., 2017; Miao et al., 2017). Overall, the simulated results of our model are in good agreement with the reported experimental findings.

**Table 1:**
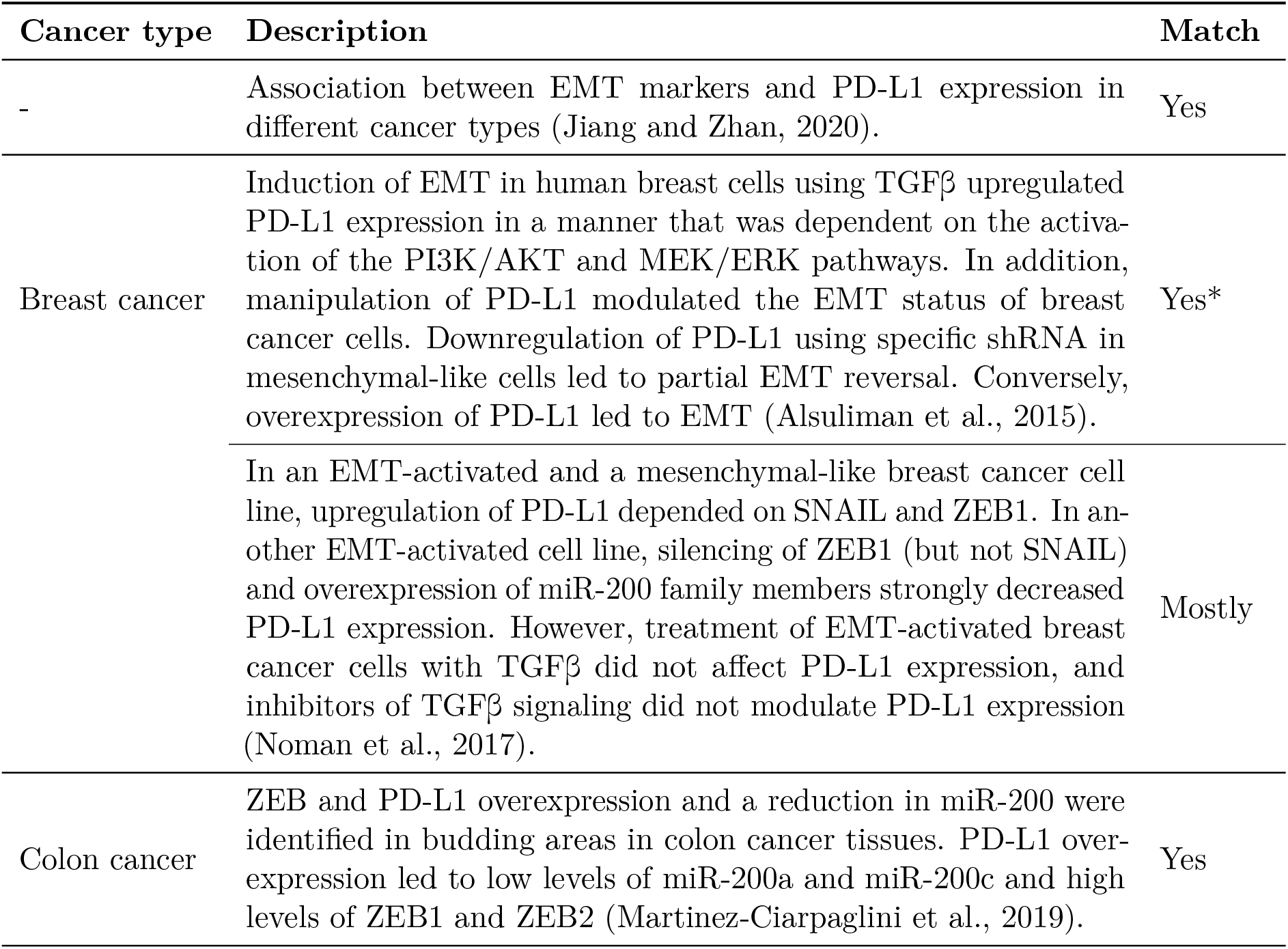
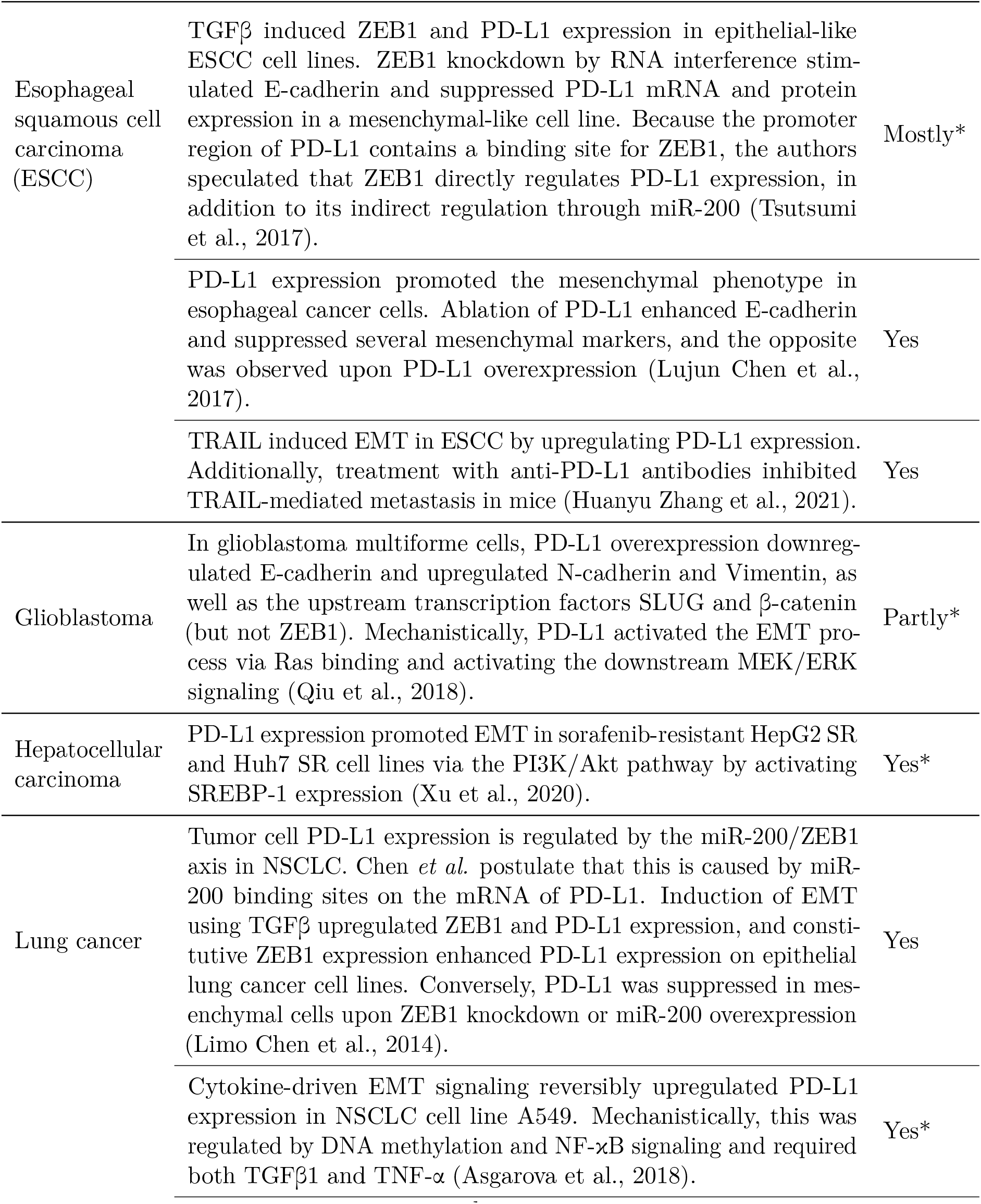
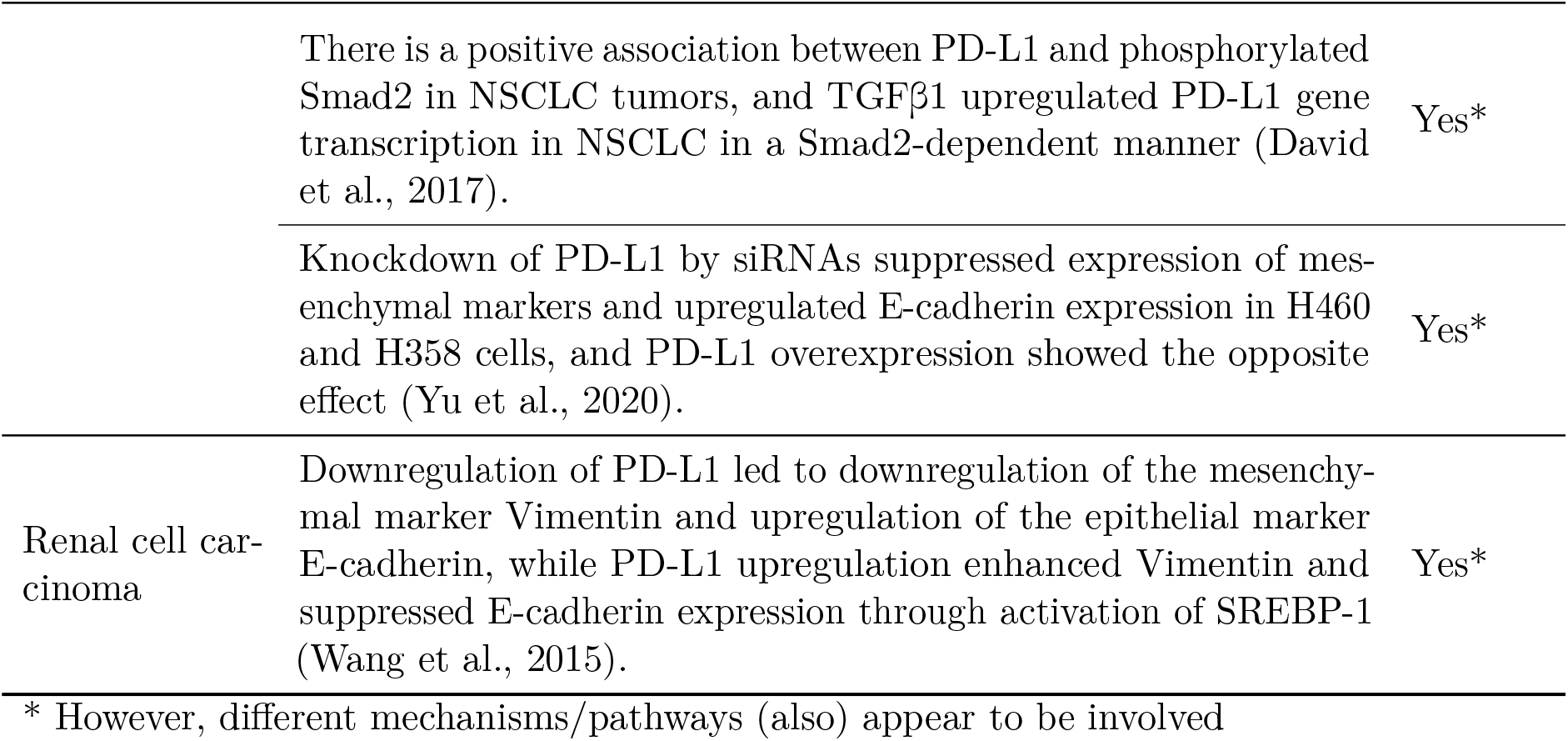
Summary of experimental reports regarding the crosstalk between EMT and PD-L1 and their consistency with our ODE model.

| Cancer type | Description | Match |
| --- | --- | --- |
| - | Association between EMT markers and PD-L1 expression in different cancer types (Jiang and Zhan, 2020). | Yes |
| Breast cancer | Induction of EMT in human breast cells using TGFβ upregulated PD-L1 expression in a manner that was dependent on the activation of the PI3K/AKT and MEK/ERK pathways. In addition, manipulation of PD-L1 modulated the EMT status of breast cancer cells. Downregulation of PD-L1 using specific shRNA in mesenchymal-like cells led to partial EMT reversal. Conversely, overexpression of PD-L1 led to EMT (Alsuliman et al., 2015). | Yes* |
| | In an EMT-activated and a mesenchymal-like breast cancer cell line, upregulation of PD-L1 depended on SNAIL and ZEB1. In another EMT-activated cell line, silencing of ZEB1 (but not SNAIL) and overexpression of miR-200 family members strongly decreased PD-L1 expression. However, treatment of EMT-activated breast cancer cells with TGFβ did not affect PD-L1 expression, and inhibitors of TGFβ signaling did not modulate PD-L1 expression (Noman et al., 2017). | Mostly |
| Colon cancer | ZEB and PD-L1 overexpression and a reduction in miR-200 were identified in budding areas in colon cancer tissues. PD-L1 overexpression led to low levels of miR-200a and miR-200c and high levels of ZEB1 and ZEB2 (Martinez-Ciarpaglini et al., 2019). | Yes |
| Esophageal squamous cell carcinoma (ESCC) | TGFβ induced ZEB1 and PD-L1 expression in epithelial-like ESCC cell lines. ZEB1 knockdown by RNA interference stimulated E-cadherin and suppressed PD-L1 mRNA and protein expression in a mesenchymal-like cell line. Because the promoter region of PD-L1 contains a binding site for ZEB1, the authors speculated that ZEB1 directly regulates PD-L1 expression, in addition to its indirect regulation through miR-200 (Tsutsumi et al., 2017). | Mostly* |
|  | PD-L1 expression promoted the mesenchymal phenotype in esophageal cancer cells. Ablation of PD-L1 enhanced E-cadherin and suppressed several mesenchymal markers, and the opposite was observed upon PD-L1 overexpression (Lujun Chen et al., 2017). | Yes |
|  | TRAIL induced EMT in ESCC by upregulating PD-L1 expression. Additionally, treatment with anti-PD-L1 antibodies inhibited TRAIL-mediated metastasis in mice (Huanyu Zhang et al., 2021). | Yes |
| Glioblastoma | In glioblastoma multiforme cells, PD-L1 overexpression downregulated E-cadherin and upregulated N-cadherin and Vimentin, as well as the upstream transcription factors SLUG and β-catenin (but not ZEB1). Mechanistically, PD-L1 activated the EMT process via Ras binding and activating the downstream MEK/ERK signaling (Qiu et al., 2018). | Partly* |
| Hepatocellular carcinoma | PD-L1 expression promoted EMT in sorafenib-resistant HepG2 SR and Huh7 SR cell lines via the PI3K/Akt pathway by activating SREBP-1 expression (Xu et al., 2020). | Yes* |
| Lung cancer | Tumor cell PD-L1 expression is regulated by the miR-200/ZEB1 axis in NSCLC. Chen *et al.* postulate that this is caused by miR-200 binding sites on the mRNA of PD-L1. Induction of EMT using TGFβ upregulated ZEB1 and PD-L1 expression, and constitutive ZEB1 expression enhanced PD-L1 expression on epithelial lung cancer cell lines. Conversely, PD-L1 was suppressed in mesenchymal cells upon ZEB1 knockdown or miR-200 overexpression (Limo Chen et al., 2014). | Yes |
| | Cytokine-driven EMT signaling reversibly upregulated PD-L1 expression in NSCLC cell line A549. Mechanistically, this was regulated by DNA methylation and NF-κB signaling and required both TGFβ1 and TNF-α (Asgarova et al., 2018). | Yes* |
| | There is a positive association between PD-L1 and phosphorylated Smad2 in NSCLC tumors, and TGFβ1 upregulated PD-L1 gene transcription in NSCLC in a Smad2-dependent manner (David et al., 2017). | Yes* |
|  | Knockdown of PD-L1 by siRNAs suppressed expression of mesenchymal markers and upregulated E-cadherin expression in H460 and H358 cells, and PD-L1 overexpression showed the opposite effect (Yu et al., 2020). | Yes* |
| Renal cell carcinoma | Downregulation of PD-L1 led to downregulation of the mesenchymal marker Vimentin and upregulation of the epithelial marker E-cadherin, while PD-L1 upregulation enhanced Vimentin and suppressed E-cadherin expression through activation of SREBP-1 (Wang et al., 2015). | Yes* |

\* However, different mechanisms/pathways (also) appear to be involved

Although findings of our model and experimental data are in good agreement, some results are partially conflicting: Firstly, Noman et al. (2017) report that while PD-L1 expression in EMT-activated breast cancer cell lines is regulated via the SNAIL1/ZEB1/miR-200 axis, treatment of MCF7-2101 cells with TGFβ or inhibition of TGFβ in MCF sh-WISP2 cells did not modulate PD-L1 expression. However, the MCF7-2101 cell line already has a mesenchymal phenotype, evident from their Vimentin expression and lack of E-cadherin; hence additional TGFβ is not expected to further increase PD-L1 expression by triggering EMT. Moreover, although the TGFβ signaling inhibitors repressed SMAD2 activation in MCF7 sh-WISP2 cells, phosphorylated SMAD2/3 is only essential for SNAIL1 activation and not for the persistence of SNAIL1 expression (J. Zhang, Tian, Y.-J. Chen, et al., 2018). Thus, the performed experiments cannot rule out that TGFβ signaling modulates PD-L1 expression in these cell lines. Secondly, PD-L1 expression in certain cell lines is reportedly affected by silencing of ZEB1 but not of SNAIL1 (Noman et al., 2017; Tsutsumi et al., 2017). Although SNAIL1 drives ZEB1 expression in the EMT core regulatory network model we used, this apparently contradictory finding can be reconciled by appreciating that the EMT transcriptional response is highly context-specific (Nieto, 2017; Cook and Vanderhyden, 2020). Indeed, sustained expression of ZEB1 can be achieved by various EMT regulators and even by ZEB1 itself, resulting in an irreversible switch to the mesenchymal phenotype (Gregory et al., 2011; J. Zhang, Tian, Hang Zhang, et al., 2014; W. Jia et al., 2019). Thus, SNAIL1 silencing may be insufficient to affect PD-L1 in settings where ZEB1 is maintained via other means.

Even though partially conflicting findings between our model and published data can be reconciled, it is also likely that additional mechanisms by which PD-L1 and EMT mutually influence each other play a role depending on the studied cell line or cancer. For instance, PD-L1 expression can induce EMT by activating the TF SREBP-1c in hepatocellular and renal cell carcinoma (Wang et al., 2015; Xu et al., 2020), ZEB1 can bind and silence IRF1, a TF of PD-L1 (Jun Yang et al., 2017), and activity of the CMTM family, which stabilizes PD-L1, correlates with EMT-TFs such as SLUG (Jie Wu, 2020; Li et al., 2021).

In the related mathematical model presented by Sahoo et al. (2021), an indirect feedback mechanism of PD-L1-mediated E-cadherin inhibition was included. Results of their model agree qualitatively with those of our model in terms of the prediction that hybrid E/M cells are expected to be PD-L1 positive and hence immune-evasive. However, the predicted PD-L1 expression levels for the different EMT phenotypes differ substantially: Sahoo et al. predict that the hybrid and mesenchymal phenotypes have almost equally high PD-L1 expression, whereas we predict that hybrid cells are closer to epithelial cells with respect to PD-L1 expression. They substantiated that PD-L1 expression increases with EMT scores by an analysis of gene expression in pan-cancer datasets and analysis of responses in several cell lines. For some of these data, intermediate EMT scores (interpreted as hybrid cells) have the tendency to only moderately increase the PD-L1 expression (e.g., Fig. 1E-F and 5E Sahoo et al., 2021), which is more in line with our model predictions. Further research will be required to unravel this complicated interplay between factors affecting PD-L1 expression and EMT. Our model forms a basis to include such factors.

A limitation of our model is its relatively crude description of IFNγ-induced PD-L1 expression. To keep the model simple, we used a steady-state approximation of the simplified model by Quaiser, Dittrich, et al. (2011), which only describes the first 15 minutes of JAK–STAT signaling before transcriptional feedback occurs. Nevertheless, our model’s predicted IRF1 temporal dynamics roughly agree with the JAK–STAT model by Rateitschak et al. (cf. Fig. 4A, 2010). Comparison of these results with those of a highly simplified IFNγ-driven PD-L1 expression model showed that the role of JAK–STAT signaling primarily creates a “switchlike” response to IFNγ (cf. Figs. S2B and 2B).

The predicted decrease of SNAIL1 levels required for EMT (as indicated by the leftward shift in the bifurcation diagrams in Figs. 2C and 2D) is noteworthy in itself; whereas several studies report on the modulation of PD-L1 expression as a result of EMP, the mechanism by which PD-L1 could modulate EMT remains unclear (Jiang and Zhan, 2020). Here we show that merely the degradation of the PD-L1-miR-200 complex could be affecting miR-200 levels sufficiently to affect EMT, without the need for additional mechanisms.

The results reported here may have diagnostic and therapeutic implications. PD-L1 has been proposed as a biomarker to predict the efficacy of PD-1/PD-L1 blockade therapy (Ren et al., 2020), yet the multifactorial mechanisms underlying PD-L1 membrane expression complicate its use as an exclusive biomarker (Patel and Kurzrock, 2015). For example, oncogene-driven PD-L1 expression caused by EMT, is constitutive and diffuse, and distinct from inflammation-driven PD-L1 expression, which may occur more focally and during a limited time window. The latter is often associated with the presence of an immune infiltrate, while the former is associated with a lack thereof (Patel and Kurzrock, 2015), and it has been proposed that this combination of PD-L1 expression and the presence of immune infiltrate affects the response to PD-1/PD-L1 blockade therapy (Lai et al., 2018). Our findings highlight how oncogenic and inflammation-driven PD-L1 expression might interact and give rise to strongly increased PD-L1 levels. PD-L1 on tumor cells is an important contributor to immune evasion through inhibition of CD8+ T cell cytotoxicity (Juneja et al., 2017). Consistently, in metastatic urothelial cancer, lack of response to anti-PD-L1 treatment occurred particularly in patients with tumors that showed exclusion of CD8+ T cells (Mariathasan et al., 2018). Combined treatment with TGFβ-blocking and anti-PD-L1 antibodies invoked anti-tumor immunity and tumor regression by facilitating T cell infiltration, which could be due to EMT-independent TGFβ signaling (Gunderson et al., 2020), but possibly also via the EMT-dependent path we have presented here. Another opportunity could be the use of therapeutic siRNA (Hu et al., 2020): In contrast to anti-PD-L1 antibodies, siRNA affects PD-L1 mRNA, which, according to our model, should free up miR-200 for its anti-EMT effect, a prediction that is supported by (partial) EMT reversal following downregulation of PD-L1 using siRNA/shRNA (Alsuliman et al., 2015; Yu et al., 2020).

In conclusion, we developed a mathematical model that describes the crosstalk between IFNγ-induced PD-L1 and EMT, which is in good agreement with experimental findings. Additionally, our model sheds light on potential mechanisms behind EMT-mediated immune evasion; and primary, adaptive, or acquired resistance to immunotherapy. Our (simplified) model can serve as a starting point to explore additional EMP and immune crosstalk mechanisms. In particular, we propose embedding the presented model in a multi-scale model to explicitly describe the local effects of the adaptive immune response and the effects of TGFβ on the tumor microenvironment. Improved understanding of the interaction between the immune response and EMP is indispensable for developing better diagnostic and therapeutic options for cancer patients.

## Materials & Methods

### Simplified Ternary Chimera Switch model

The simplified TCS model (Jolly, Tripathi, D. Jia, et al., 2016) is built on the theoretical framework for microRNA-TF chimera toggle-switches defined in M. Lu, Jolly, Gomoto, et al. (2013). See Supplementary Information for the model definition, theoretical framework, and used parameters.

### IFNγ–PD-L1 model

The IFNγ–PD-L1 model consists of two parts: (IFNγ–)JAK–STAT signaling, for which we use a steady-state approximation of an existing model, and STAT–PD-L1 regulation which we developed following the theoretical framework underpinning the simplified TCS model. We discuss these parts in more detail below.

### (IFNγ–)JAK–STAT

For the IFNγ–PD-L1 model, we use the JAK–STAT model by (Quaiser, Dittrich, et al., 2011) (see Supplementary Information). Because the dynamics of STAT–PD-L1 and EMT signaling in the TCS model take place on a much longer time scale than those of the JAK–STAT model (see Fig. 1C), we simplified our IFNγ–PD-L1 model by assuming the JAK–STAT model is in quasi-steady state; we fitted its steady-state by a Gompertz function (see Fig. S5) and inserted this approximate relationship between IFNγ and STAT in our STAT–PD-L1 model.

### STAT–PD-L1

The STAT–PD-L1 submodel (see also Fig. S1A) consists of the following ordinary differential equations (ODEs):

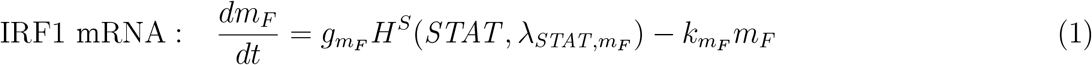

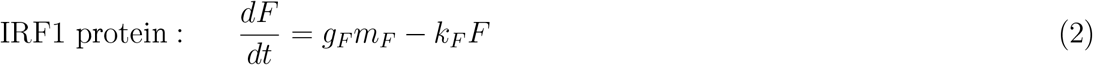

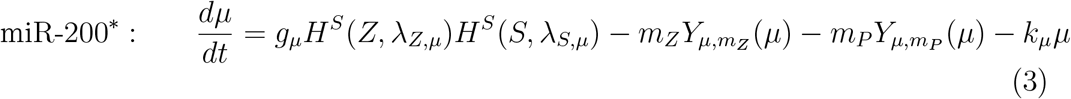

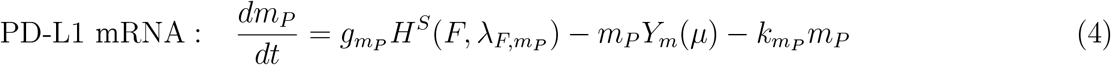

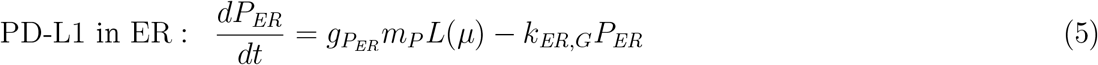

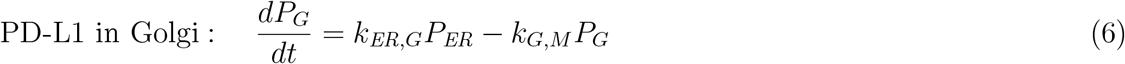

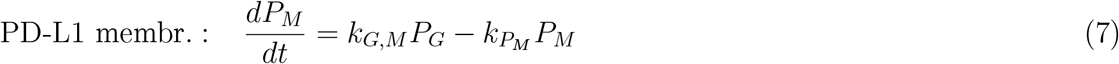

This model uses the abundance of STAT (in molecules) as input, and includes IRF1 and PD-L1 using appropriate TF-TF and miR-TF dynamics (*H*^*S*^, *L*, and *Y* functions) from the theoretical framework by M. Lu, Jolly, Gomoto, et al. (2013) (see Supplementary Information) in anticipation of the eventual connection to miR-200 (*μ*) in our combined model (see below). To accommodate a low number of binding sites for miR-200 on the mRNA of PD-L1 (Limo Chen et al., 2014), we adapted the parameters for the *L*(*μ*), *Y*_*m*_(*μ*), and 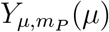 from Table S1 (*n* = 6) to the values as shown in Table 2 (*n* = 2). We assumed *μ*_0_ = 10^4^ molecules to be the same as in Jolly, Tripathi, D. Jia, et al. (2016). Transport of PD-L1 through various cellular compartments is modeled with a constant rate from compartment to compartment, with rates assumed to be similar to Lippincott-Schwartz, Roberts, and Hirschberg (2000). Model parameters are provided in Table 3; some parameters were considered to be similar to those of the simplified TCS model.

**Table 2:** Rates used in the *L*(*μ*), *Y*_*m*_(*μ*), and 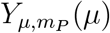 functions in the STAT-PD-L1 model for *n* = 2.

| $n$ (# of miRNA binding sites) | 0 | 1 | 2 |
| --- | --- | --- | --- |
| $l_i$ [h<sup>-1</sup>] | 1 | 0.3 | 0.05 |
| $\gamma_{mi}$ [h<sup>-1</sup>] | 0 | 0.2 | 1 |
| $\gamma_{\mu i}$ [h<sup>-1</sup>] | 0 | 0.05 | 0.5 |

**Table 3:** Variables and parameters used for the STAT–PD-L1 model. The top panel shows variable names and production and degradation rates; the bottom panel shows parameters for the shifted Hill functions of the interactions. Degradation rates *k* are in h^−1^, production rates *g* in molecules h^−1^, and thresholds 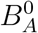 in molecules.

| | | | Prod. rate $g$ | Degr. rate $k$ | |
| --- | --- | --- | --- | --- | --- |
| IRF1 mRNA | $m_F$ | $g_{m_F}$ | 30 | $k_{m_F}$ | 0.5 |
| IRF1 protein | $F$ | $g_F$ | 100 | $k_F$ | 0.1 |
| PD-L1 mRNA | $m_P$ | $g_{m_P}$ | 30 | $k_{m_Z}$ | 0.5 |
| PD-L1 in ER | $P_{ER}$ | $g_{P_{ER}}$ | 100 | $k_{ER,G}$ | 1.68 |
| PD-L1 in Golgi Complex | $P_G$ | | | $k_{G,M}$ | 1.8 |
| PD-L1 on membrane | $P_M$ | | | $k_{P_M}$ | 0.15 |

| | | Threshold $B_A^0$ | Hill coefficient $n_{BA}$ | | Max. fold change $\lambda_{BA}$ | |
| --- | --- | --- | --- | --- | --- | --- |
| Act. $m_F$ by $STAT$ | $STAT_{m_F}^0$ | $2 \times 10^6$ | $n_{STAT, m_F}$ | 10 | $\lambda_{STAT, m_F}$ | 10 |
| Act. $m_P$ by $F$ | $F_{m_P}^0$ | $10^5$ | $n_{F, m_P}$ | 3 | $\lambda_{F, m_P}$ | 10 |

### Combined model

To create our combined model, we connected the IFNγ-induced JAK–STAT signaling model and simplified TCS model to the central STAT–PD-L1 model (see Fig. 1B). Since the JAK–STAT model has [STAT1p_2] as output in nm we need to convert to number of molecules to use STAT1p_2 as input in the STAT–PD-L1 model. As in Jolly, Tripathi, D. Jia, et al. (2016), we use a cell volume of 10 000 μm^3^, such that 1 nm amounts to roughly 6020 molecules (6.02 × 10^23^ ·10^−9^ ·10000 × (10^−5^)^3^). Note that this cell volume is on the high side, as typical animal cells are 10–20 μm in diameter (∼ 500–4000 μm^3^) (Alberts et al., 2015, p. 529). However, because of our IFNγ–STAT steady-state approximation, a decrease in cell volume by a factor 10 can be compensated by multiplying the inducing IFNγ signal also by a factor 10 to yield the same amount of STAT, and hence, the same model result. Additionally, to connect the STAT–PD-L1 model to the simplified TCS using miR-200, equation miR-200* (Eq. (3)) is used instead of the equation for miR-200 in the simplified TCS model to include the interaction with PD-L1, and 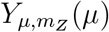 in the updated equation corresponds to *Y*_*μ*_(*μ*) in the simplified TCS model (see Supplementary Information). Table 4 shows all model components and their units in our combined model.

**Table 4:** List of regulators in the combined model. Starred (*) regulators IFNγ and SNAIL1 are the two model inputs.

| Regulator | Symbol | Units |
| --- | --- | --- |
| *extra-cellular IFNγ | $[IFN\gamma]$ | nM |
| [STAT1p_2] | $x_{10}$ | nM |
| STAT1p_2 | $STAT$ | # molecules |
| IRF1 mRNA | $m_F$ | # molecules |
| IRF1 protein | $F$ | # molecules |
| PD-L1 mRNA | $m_P$ | # molecules |
| PD-L1 in Endoplasmatic Reticulum | $P_{ER}$ | # molecules |
| PD-L1 in Golgi Complex | $P_G$ | # molecules |
| PD-L1 on cell membrane | $P_M$ | # molecules |
| miR-200 | $\mu$ | # molecules |
| ZEB1 mRNA | $m_Z$ | # molecules |
| ZEB1 protein | $Z$ | # molecules |
| *SNAIL1 protein | $S$ | # molecules |

### Simulation & Analysis

For model simulations, we used COPASI (COmplex PAthway SImulator) (RRID:SCR_014260) (Hoops et al., 2006), and model files are included in the Supplementary Information.

Analysis in R (R Project for Statistical Computing, RRID:SCR_001905) (R Core Team, 2018) was performed with RStudio (RStudio, RRID:SCR_000432) (RStudio Team, 2016) and the tidyverse (Wickham, 2017) packages.

## Supporting information

COPASI model files

## Acknowledgments

This work was supported by a Vidi grant from the Netherlands Organization for Scientific Research (NWO; grant 864.12.013 to JBB).

Figure 1A was created in BioRender.com.

## Supplementary Information

### Supplementary Methods

#### Simplified Ternary Chimera Switch model

The TCS model (M. Lu, Jolly, Levine, et al., 2013) is built on the theoretical framework for microRNA-TF chimera toggle-switches defined in M. Lu, Jolly, Gomoto, et al. (2013). In this framework, activation and inhibition of TF B on TF A is defined by a shifted Hill function

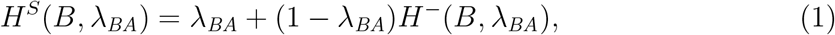

where 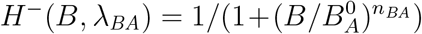 is an inhibitory Hill function, 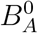 is the threshold level for *B* at which inhibition is half-maximal, *n*_*BA*_ is the Hill coefficient (usually associated with the number of binding spots on the promoter), and *λ*_*BA*_ is the maximum fold change of *A* caused by *B* (when B is an inhibitor, this implies that 0 ≤ *λ*_*BA*_ *<* 1). The miRNA-affected translation rate of mRNA is given by

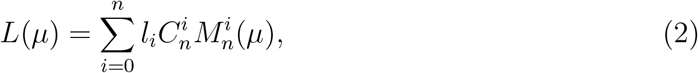

such that the total translation is *L*_tot_ = *L*(*μ*)*m*_0_, where *m*_0_ is the total concentration of mRNA. The miRNA-assisted mRNA degradation rate is given by

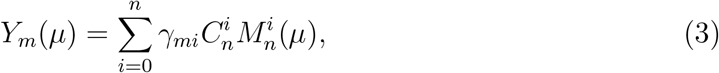

and the miRNA-assisted miRNA degradation rate by

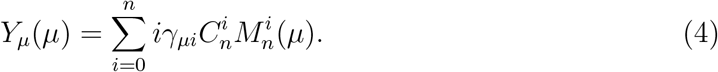

In these equations

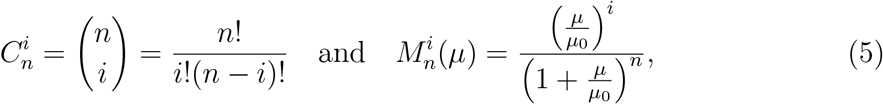

and

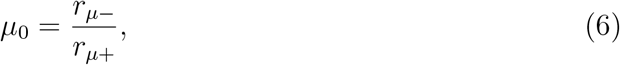

where *μ* is the amount of miRNA, *r*_*μ*+_ and *r*_*μ*−_ the binding and unbinding rate miRNA to mRNA, *n* is the number of binding sites, *l*_*i*_ is the translation rate of an mRNA when bound to *i* miRNAs, *γ*_*mi*_ is the individual degradation rate of mRNA bound to *i* molecules of miRNA, and *γ*_*μi*_ is the individual degradation rate for a miRNA molecule (see SI M. Lu, Jolly, Levine, et al., 2013, for derivation and more details).

For the simplified TCS model, with SNAIL as input (M. Lu, Jolly, Levine, et al., 2013; Jolly, Tripathi, D. Jia, et al., 2016), the parameters used in *L*(*μ*), *Y*_*m*_(*μ*), and *Y*_*μ*_(*μ*) functions are shown in Table S1.

**Table S1:** Rates used in the *L*(*μ*), *Y*_*m*_(*μ*), and *Y*_*μ*_(*μ*) functions in the simplified TCS model. Other parameters are fixed at *μ*_0_ = 10^4^ molecules and *n* = 6 (Jolly, Tripathi, D. Jia, et al., 2016).

| $n$ (# of miRNA binding sites) | 0 | 1 | 2 | 3 | 4 | 5 | 6 |
| --- | --- | --- | --- | --- | --- | --- | --- |
| $l_i$ [$\text{h}^{-1}$] | 1 | 0.6 | 0.3 | 0.1 | 0.05 | 0.05 | 0.05 |
| $\gamma_{mi}$ [$\text{h}^{-1}$] | 0 | 0.04 | 0.2 | 1 | 1 | 1 | 1 |
| $\gamma_{\mu i}$ [$\text{h}^{-1}$] | 0 | 0.005 | 0.05 | 0.5 | 0.5 | 0.5 | 0.5 |

Using this theoretical framework we can now write the simplified TCS model as

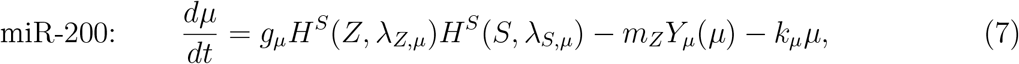

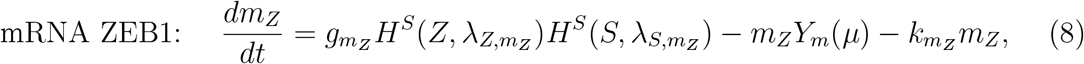

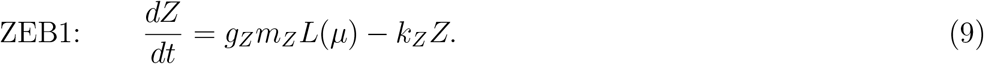

Table S2 lists the variables and parameters used in the simplified TCS model.

**Table S2:** Parameters used for the TCS model. Top panel shows production and degradation rates, bottom panel shows parameters for the shifted Hill functions of the interactions. Degradation rates *k* and *l* are in h^−1^, production rates *g* in molecules h^−1^, and thresholds 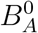 in molecules (Jolly, Tripathi, D. Jia, et al., 2016).

| | | Prod. rate $g$ | | Degr. rate $k$ | |
| --- | --- | --- | --- | --- | --- |
| miR-200 | $\mu$ | $g_\mu$ | 2100 | $k_\mu$ | 0.05 |
| mRNA ZEB1 | $m_Z$ | $g_{m_Z}$ | 11 | $k_{m_Z}$ | 0.5 |
| ZEB1 | $Z$ | $g_Z$ | 100 | $k_Z$ | 0.1 |

| | Threshold $B_A^0$ | Hill coefficient $n_{BA}$ | Max. fold change $\lambda_{BA}$ |
| --- | --- | --- | --- |
| Inh. $\mu$ by $Z$ | $Z_\mu^0$ $2.2 \times 10^5$ | $n_{Z,\mu}$ 3 | $\lambda_{Z,\mu}$ 0.1 |
| Inh. $\mu$ by $S$ | $S_\mu^0$ $1.8 \times 10^5$ | $n_{S,\mu}$ 2 | $\lambda_{S,\mu}$ 0.1 |
| Act. $m_Z$ by $Z$ | $Z_{m_Z}^0$ $2.5 \times 10^4$ | $n_{Z,m_Z}$ 2 | $\lambda_{Z,m_Z}$ 7.5 |
| Act. $m_Z$ by $S$ | $S_{m_Z}^0$ $1.8 \times 10^5$ | $n_{S,m_Z}$ 2 | $\lambda_{S,m_Z}$ 10 |

#### JAK–STAT model

To model JAK–STAT signaling, we use model *M* ^0^ by Quaiser, Dittrich, et al. (2011), which is a truncated model from Yamada et al. (2003), developed to resolve non-identifiability in the non-truncated model (Quaiser and Mönnigmann, 2009). The truncated model focuses on the first 15 minutes of JAK–STAT signaling, where transcriptional feedback does not occur yet. The model is given by:

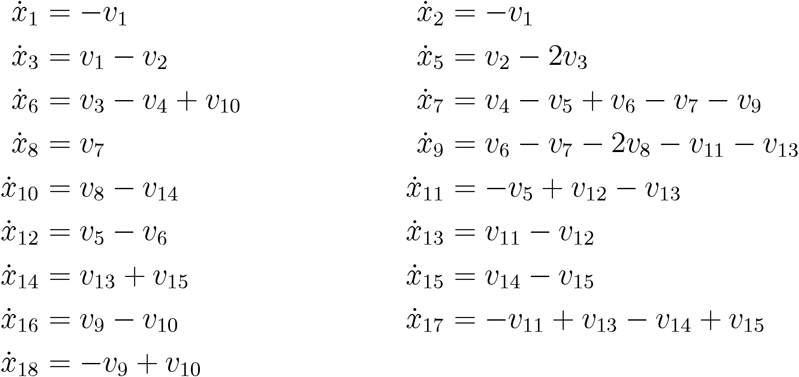

where the *v* parameters are defined as follows:

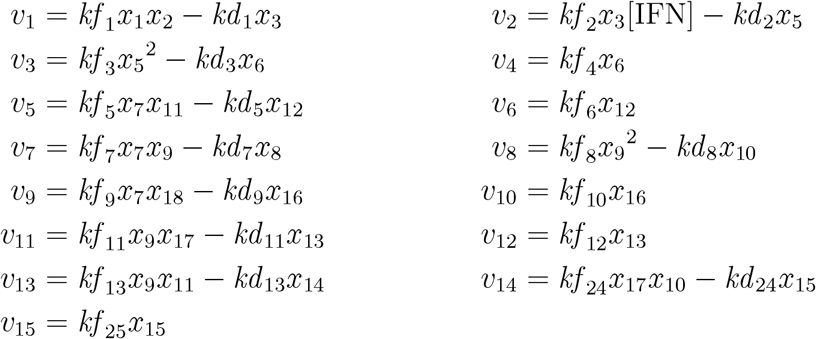

In this model, *x*_4_ represents the input concentration of IFNγ, and *x*_10_ the output concentration of STAT1_p2 in nm. Table S3 contains the initial conditions and parameters (Quaiser, Dittrich, et al., 2011).

Because this model describes a much shorter time scale (∼ 15 min) than that of the PD-L1 dynamics (∼ 1 d), and EMT process (∼ 20 d), we essentially used the quasi-steady-state assumption for the JAK–STAT model; to keep the full model relatively simple, we first obtained the numeric steady-steady of [STAT1p_2] for different values of IFNγ using COPASI, then we approximated the dependence on this quasi-steady state by fitting it to a three-parameter Gompertz function (Fig. S5), which can be parameterized as follows:

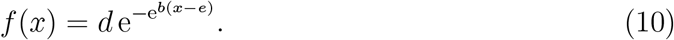

Fitting was done by the G.3 self-starter in the drc R package (Ritz et al., 2015), yielding parameter values *d* = 414.56394, *b* = −51.52794, and *e* = 0.02833.

**Table S3:** Initial conditions and parameters for the JAK–STAT model. Obtained from Tables S1 and S2a from Quaiser, Dittrich, et al. (2011).

| Initial condition nM |  | Parameters s<sup>-1</sup> |  |  |  |
| --- | --- | --- | --- | --- | --- |
| $x_1$ | 12 | $kf_1$ | 0.1 | $kd_1$ | $0.5 \times 10^{-1}$ |
| $x_2$ | 12 | $kf_2$ | $0.2 \times 10^{-1}$ | $kd_2$ | $0.2 \times 10^{-1}$ |
| $x_3$ | 0 | $kf_3$ | $0.4 \times 10^{-1} \text{ nM}^{-1}$ | $kd_3$ | 0.2 |
| $x_5$ | 0 | $kf_4$ | $0.5 \times 10^{-2}$ | | |
| $x_6$ | 0 | $kf_5$ | $0.8 \times 10^{-2}$ | $kd_5$ | 0.8 |
| $x_7$ | 0 | $kf_6$ | 0.4 | | |
| $x_8$ | 0 | $kf_7$ | $0.5 \times 10^{-2}$ | $kd_7$ | 0.5 |
| $x_9$ | 0 | $kf_8$ | $0.2 \times 10^{-1} \text{ nM}^{-1}$ | $kd_8$ | 0.1 |
| $x_{10}$ | 0 | $kf_9$ | $0.1 \times 10^{-2}$ | $kd_9$ | 0.2 |
| $x_{11}$ | 1000 | $kf_{10}$ | $0.3 \times 10^{-2}$ | | |
| $x_{12}$ | 0 | $kf_{11}$ | $0.1 \times 10^{-2}$ | $kd_{11}$ | 0.2 |
| $x_{13}$ | 0 | $kf_{12}$ | $0.3 \times 10^{-2}$ | | |
| $x_{14}$ | 0 | $kf_{13}$ | $2 \times 10^{-7}$ | $kd_{13}$ | 0.2 |
| $x_{15}$ | 0 | $kf_{24}$ | $0.1 \times 10^{-2}$ | $kd_{24}$ | 0.2 |
| $x_{16}$ | 0 | $kf_{25}$ | $0.3 \times 10^{-2}$ | | |
| $x_{17}$ | 50 | | | | |
| $x_{18}$ | 100 | | | | |

### Figures

**Figure S1:**
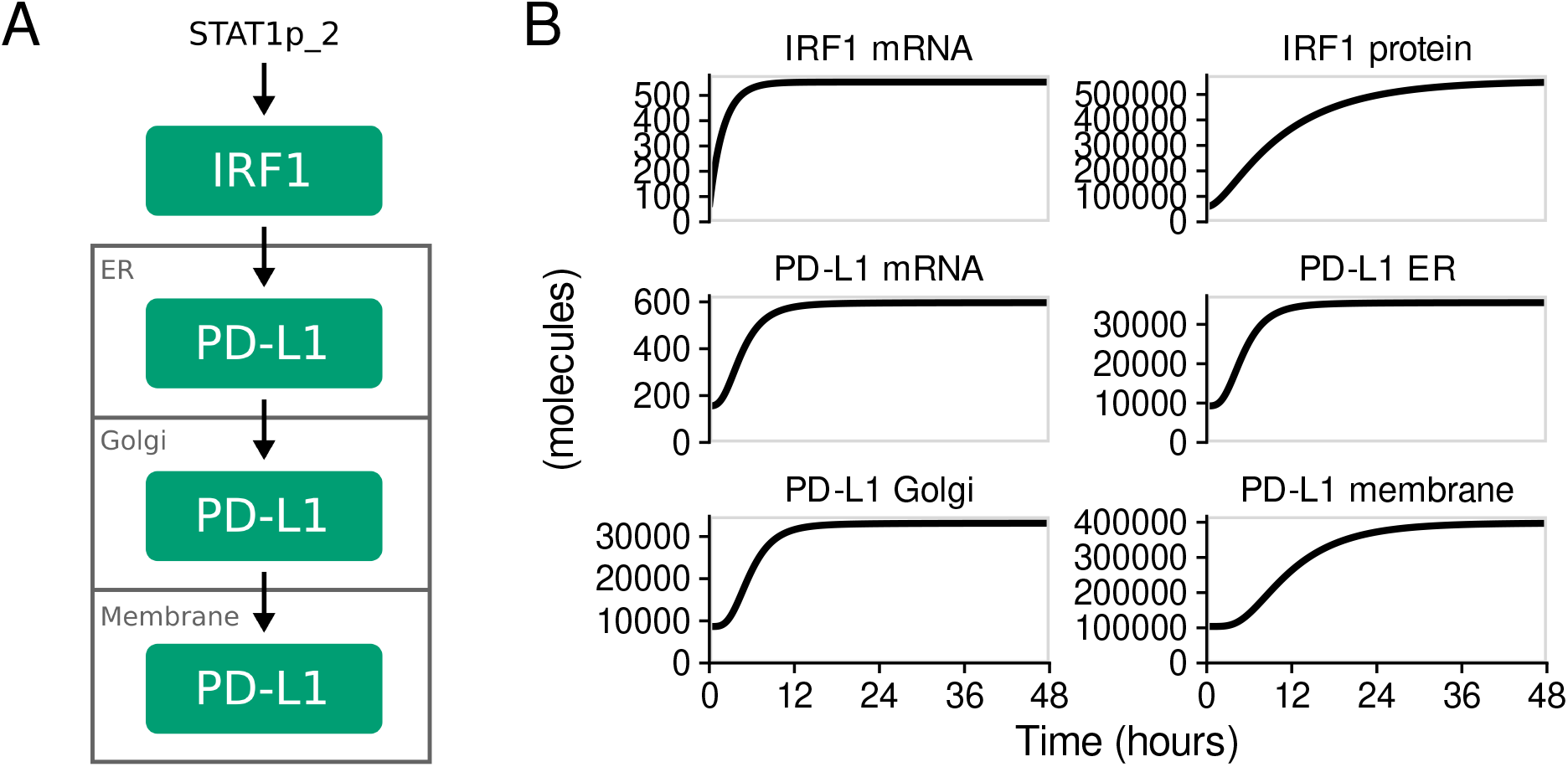
Model for STAT1-induced PD-L1 expression. (A) Schematic diagram that shows how STAT1 expression drives IRF1 and PD-L1 in the Endoplasmatic Reticilum (ER), after which PD-L1 is transported to the membrane. (B) Temporal dynamics of STAT1-induced PD-L1 expression following induction with STAT1p_2 = 2.5 × 10^6^ molecules.

**Figure S2:**
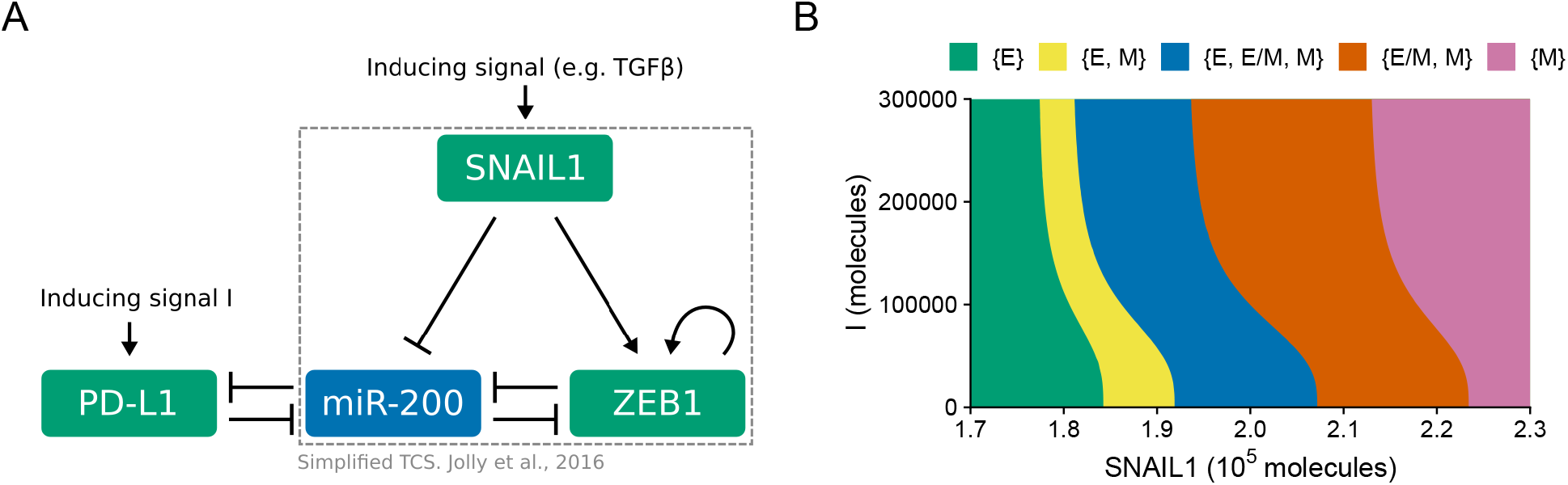
Simplified interaction model. (A) Schematic diagram of our simplified model driven by a direct inducing signal I and SNAIL1. (B) Phase diagram of our simplified model that shows the possible coexistence of the different EMT phenotypes.

**Figure S3.**
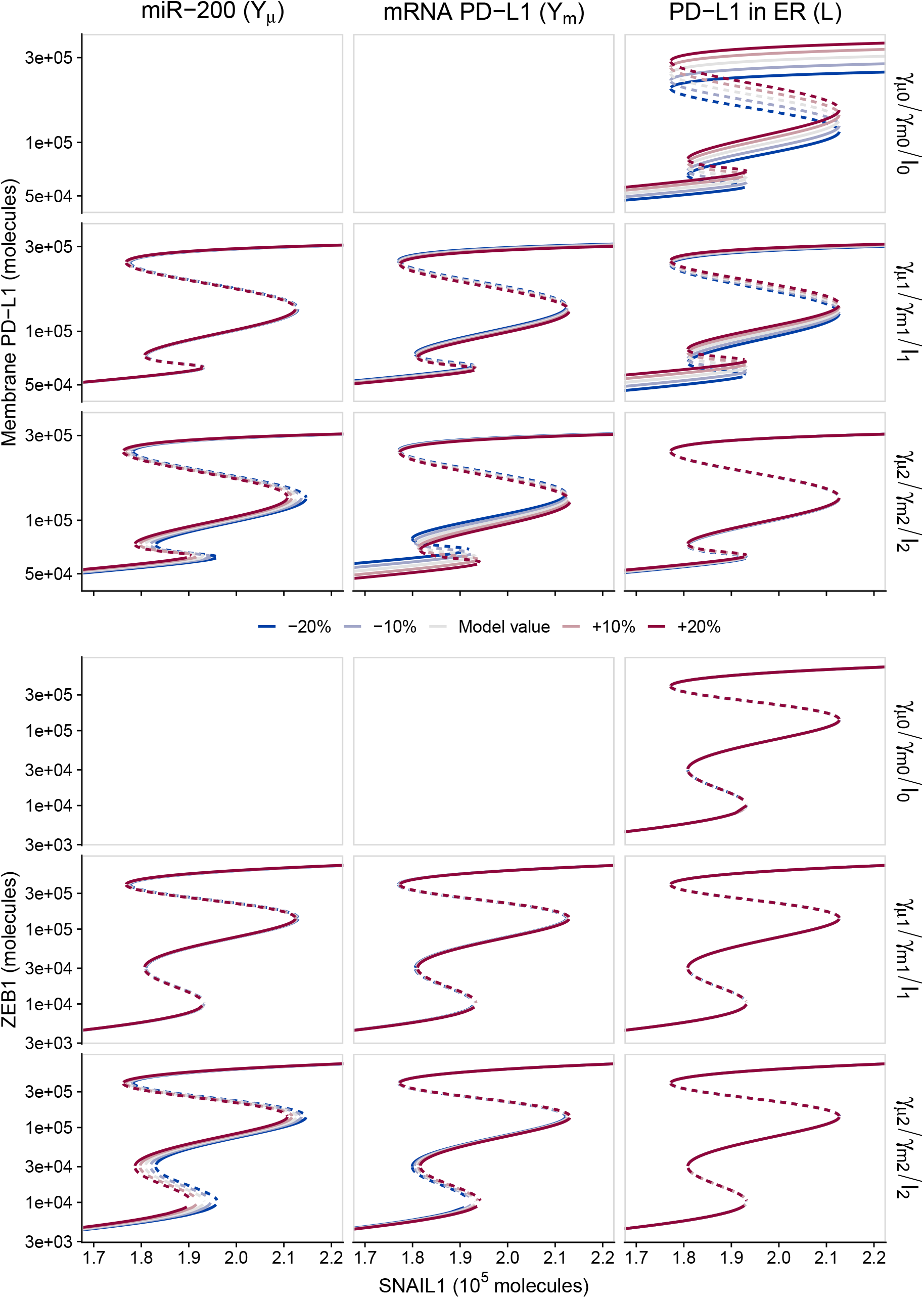
Sensitivity analysis of PD-L1 at the membrane and ZEB1 protein for parameters used in the negative-feedback loop between miR-200 and PD-L1. Modifications in the bifurcation diagram are shown for PD-L1 (top) and ZEB1 (bottom) as dependent on SNAIL1 but for fixed IFNγ = 0.1 nm. Parameters *γ*_*μi*_ (left), *γ*_*mi*_ (middle), and *l*_*i*_ (right) for the 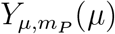, *Y*_*m*_(*μ*), and *L*(*μ*) functions (see Table 2) were varied from the default model values (black) by +10/20% (red shades) and -10/20% (blue shades).

**Figure S4:**
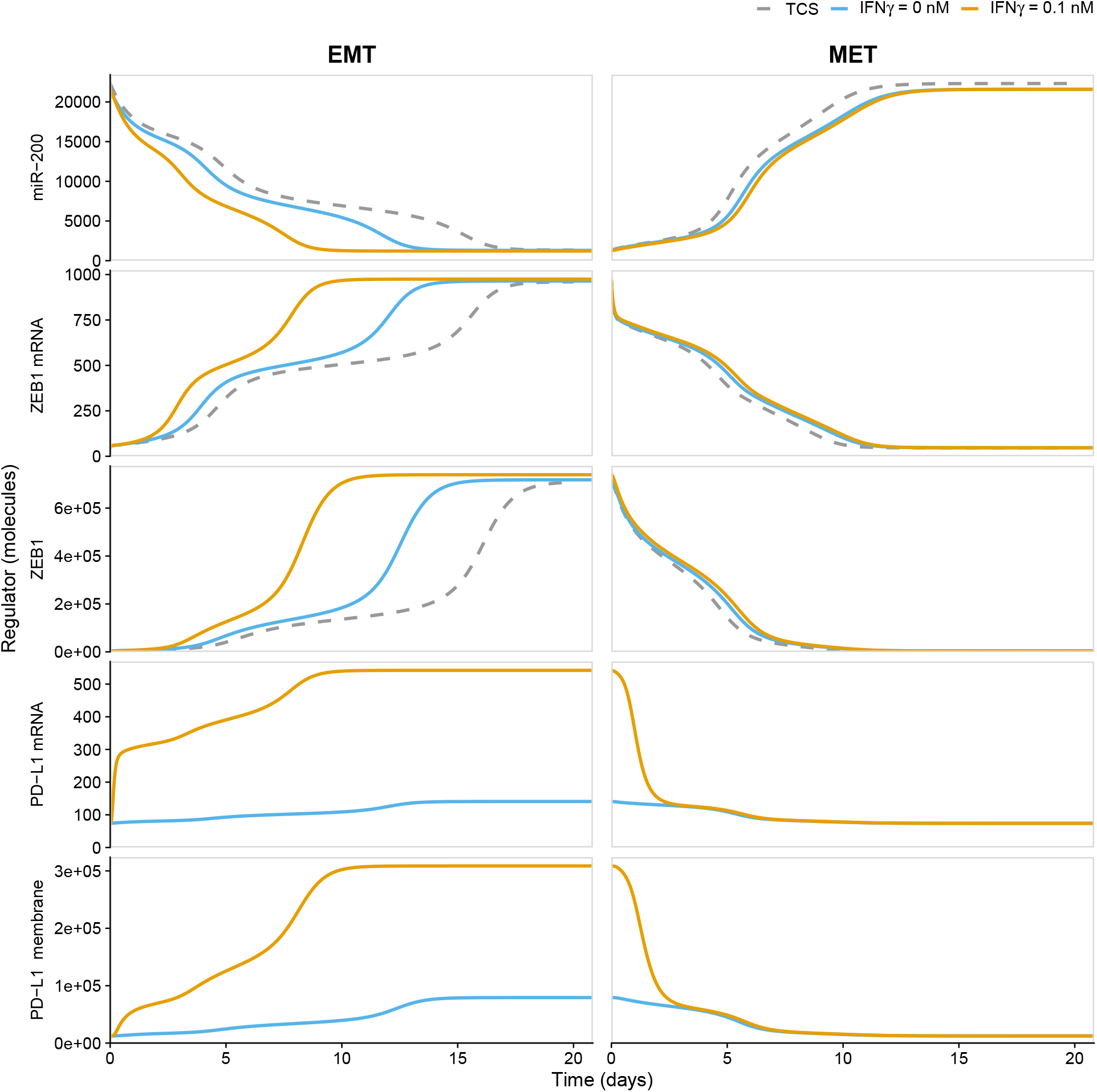
Temporal dynamics of EMP after simultaneous IFNγ and EMT induction. EMT (left) and MET (right) for the simplified TCS model (dashed gray), and our combined model (Fig. 1B). For EMT, cells in epithelial state with SNAIL1 = 1.7 × 10^5^ molecules and IFNγ = 0 nm undergo a full EMT by simultaneously increasing SNAIL1 to 2.3 × 10^5^ molecules and IFNγ to 0.1 nm (orange), or by just increasing SNAIL1 (blue, same as blue in Fig. 3 in the main text). For MET, SNAIL1 and IFNγ are decreased to 1.7 × 10^5^ molecules and 0 nm again.

**Figure S5:**
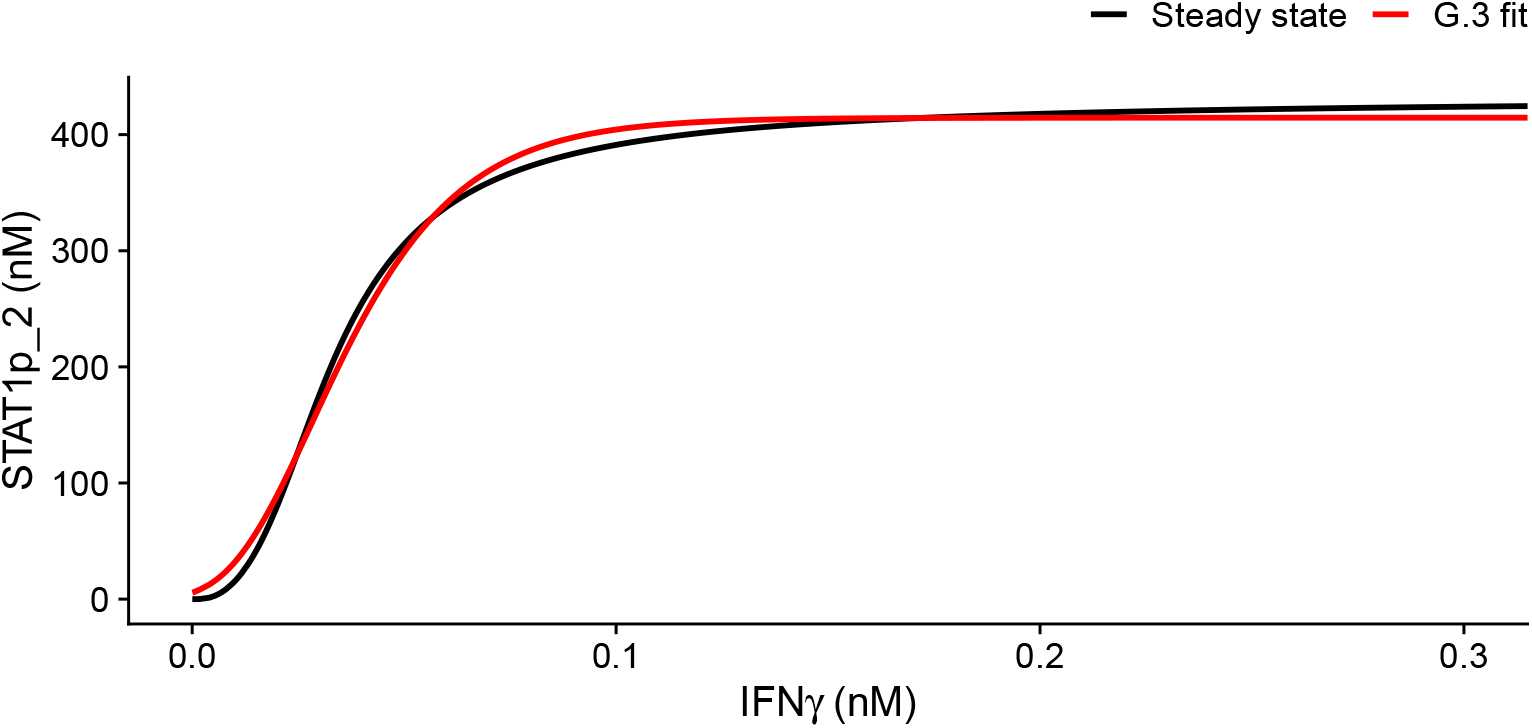
Steady-state approximation of [STATp_2] driven by IFNγ. The black line shows the simulated values of STAT in steady state, the red line shows the fit by a three-parameter Gompertz curve.

### COPASI model files

The following COPASI model files are included:

- `core_emt.cps`: The simplified TCS model as published by Jolly, Tripathi, D. Jia, et al. (2016).
- `jakstat.cps`: The JAK–STAT model as published by Quaiser, Dittrich, et al. (2011).
- `ifn_jak_stat_pdl.cps`: Our IFNγ–PD-L1 model including steady-state approximation of the JAK–STAT model.
- `full_model.cps`: Our full model combining IFNγ-induced PD-L1 production with the simplified TCS model.
- `core_emt_plus_pdl.cps`: Our simplified model where we only consider the simplified TCS model coupled to PD-L1, induced by an external signal *I*.

